# Massive export of diazotrophs across the South Pacific tropical Ocean

**DOI:** 10.1101/2021.05.07.442706

**Authors:** Sophie Bonnet, Mar Benavides, Frédéric A.C. Le Moigne, Mercedes Camps, Antoine Torremocha, Olivier Grosso, Céline Dimier, Dina Spungin, Ilana Berman-Frank, Laurence Garczarek, Francisco M. Cornejo-Castillo

## Abstract

Diazotrophs are widespread microorganisms that alleviate nitrogen limitation in 60% of our oceans, regulating marine productivity. Yet, their contribution to organic matter export has not been quantified, making an assessment of their impact on the biological carbon pump impossible. Here, we demonstrate that cyanobacterial and non-cyanobacterial diazotrophs are massively exported down to 1000 m-depth in the western subtropical South Pacific Ocean (WTSP), accounting for up to 52-100% of the total particulate nitrogen export fluxes. We further demonstrate that small size unicellular diazotrophs (UCYN, 1-8 µm) are exported more efficiently than filamentous diazotrophs (>100-1000 µm) under the form of large (>50 µm) aggregates linked by an extracellular organic matrix. Beyond the WTSP, our data are supported by analysis of the Tara Oceans metagenomes collected in other ocean basins, showing that diazotrophs are systematically detected in mesopelagic waters when present at the surface. We thus conclude that diazotrophs are key players in carbon sequestration in the ocean and need to be considered in future studies to improve the accuracy of current regional and global estimates of export.

## Introduction

Nitrogen (N) availability limits primary productivity throughout much of the surface low-latitude ocean^1^. In such nitrogen (N)-limited waters, microbial dinitrogen (N_2_) fixation by diazotrophic plankton provides the major source of new N to surface waters^2^ that maintains the ocean fertility and, on appropriate timescales, equates export production to the deep ocean^3^. However, the fate of this production remains obscure^4^. No consensus as whether it is exported to the deep ocean or stimulates remineralization in surface waters currently exists. An increasing number of studies have shown that diazotroph-derived N is quickly translocated to non-diazotrophic plankton such as diatoms^5,6^, which eventually contribute to secondary export of organic matter out of the photic zone. Yet, the direct gravitational settling of diazotrophs themselves to the deep ocean has rarely been assessed.

Diazotrophs may associate with sinking particles and contribute to direct export by different mechanisms. The most direct ones include gravitational settling of individual cells/filaments or aggregates. According to Stokes’ Law, particle sinking velocity scales with the square of particle size. Therefore, large particles should sink faster and are more likely to reach the deep ocean before being remineralized by bacteria^7^. Aggregation is thus a crucial step for the transport of small phytoplankton species which could export particulate organic carbon (POC) and N (PON) in similar proportion to their production in surface waters^8^. Diazotrophs have diverse morphologies and their size spans several orders of magnitude. Some types such as the free-living unicellular diazotrophic cyanobacteria (UCYN from groups B and C) are small (2-8 μm in diameter), while others such as *Trichodesmium sp*. are filamentous and can form large-sized colonies (>100-1000 µm). In addition, some diazotrophs live in symbiosis with calcified (UCYN-A, ∼1 μm) or silicified eukaryotes (*Richelia* sp., *Calothrix* sp., >20 µm, forming Diatom-Diazotrophic Associations or DDAs). These dense biominerals may provide ballast enhancing the downward export of these symbioses into the deep ocean. Therefore, the presence of different diazotrophs in surface waters may result in drastically different POC and PON export fluxes via the biological pump. Yet, no field observations have related individual diazotroph groups to the magnitude of downward particles fluxes to date.

Thanks to their inherently ballasted character, DDAs are well known to contribute to particulate matter export^9,10^, and are involved in seasonal peaks of POC export to the deep sea (4000 m) in the North Pacific subtropical gyre^11^. *Trichodesmium* is one of the major contributors to global N_2_ fixation^12^. However, *Trichodesmium* are thought to have a limited export capacity and to be preferentially remineralized in the surface layers due to the presence of gas vesicles providing them buoyancy^4,13,14^. Yet, some studies have reported the presence of *Trichodesmium* in sediment traps material in the Kuroshio Current^15^, the tropical North Atlantic^16^, North^17^ and Pacific Oceans^18^. Intact filaments and colonies of *Trichodesmium* sp. have also been reported as deep as 3000-4000 m in the tropical Atlantic, Pacific and Indian Oceans^16,19^, but their contribution to organic matter export has yet to be quantified.

Theoretically, UCYN may not contribute importantly to POC export fluxes either due to their small size. Yet, Berthelot et al.^20^ reported in experimental mesocosms that primary production supported by UCYN was twice as more efficient in promoting POC export than production supported by DDA. During the same study, Bonnet et al.^21^ showed that the sinking of UCYN-B and C was driven by the aggregation of small individual cells (5-6 μm) into larger aggregates (100 to >500 μm) that were eventually exported and accounted for up to ∼20% of the POC flux. Recently, Caffin et al.^18^ confirmed the presence of UCYN-B in sinking particles collected with sediment traps (330 m) throughout the western tropical South Pacific Ocean (WTSP) (10^3^ to 10^7^ *nifH* gene copies L^-1^ of trap material). Finally, Farnelid et al.^22^ observed *nifH* gene sequences in exported material (150 m) in the North Pacific subtropical gyre and found that all the diazotroph groups above as well as non-cyanobacterial diazotrophs were present in the samples.

Taken together, these studies suggest that all diazotroph groups, small or large, free-living or symbiotic, ballasted or not, have been detected below the photic layer. However, a detailed examination relating types of diazotrophs to the magnitude of downward particles fluxes and their contribution is needed to refine our understanding of their role in the magnitude and mechanisms controlling the biological carbon pump. This is even more timely as climate models predict an expansion of the oligotrophic gyres^23,24^, where diazotrophs represent a major component of the plankton biomass and sustain most new primary production^25^.

Here, we examine the species-specific fate of diazotrophs in the mesopelagic ocean. We used an innovative approach consisting of the combined deployment of surface-tethered drifting sediment traps, Marine Snow Catcher (MSC), and Bottle-net, in which we performed *nifH* sequencing and quantitative PCR on major diazotroph groups across the WTSP in parallel with biogenic element export fluxes quantification (Fig. 1a). This study builds up on prior investigations that have characterized this region as a hotspot for marine N_2_ fixation^26^. We show that all globally-significant N_2_-fixing cyanobacteria and non-cyanobacterial diazotrophs are systematically present in sinking particles down to 1000 m, where they account for a large fraction of exported PON (52-100%). Small size UCYN (1-8 µm) are exported more efficiently than large filamentous diazotrophs under the form of large (>50 µm) aggregates linked by an extracellular matrix. Globally, our analysis of the *Tara* Oceans metagenomes confirms that diazotrophs are always detected in mesopelagic waters when present in surface waters, revealing that the gravitational settling of diazotrophs is an important pathway for diazotroph-derived export production, influencing the ability of our ocean to sequester carbon.

**Figure 1.**
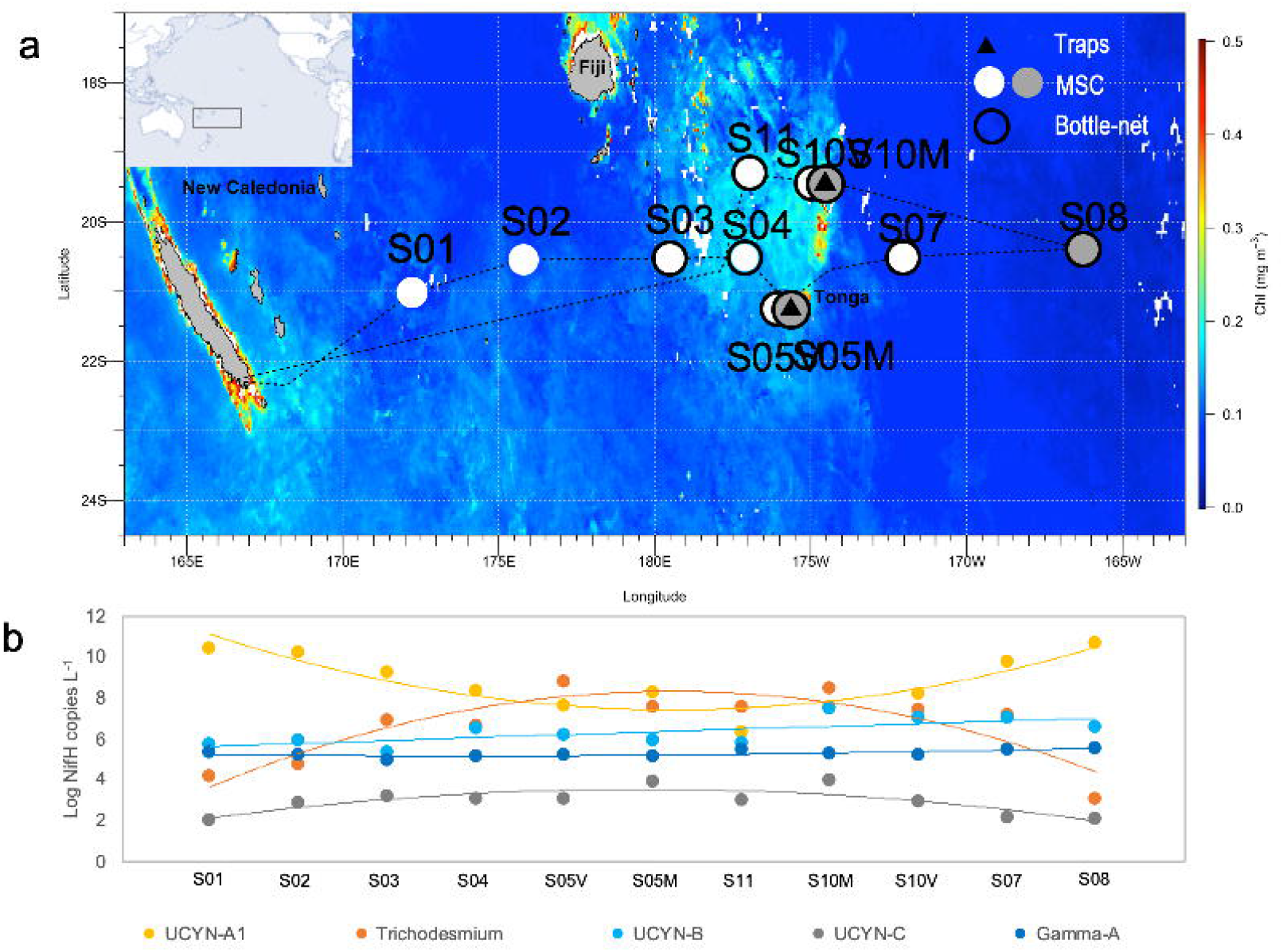
a. Surface chlorophyll MODIS composite averaged over the time period corresponding to the TONGA cruise (1 November-6 December 2019) at a resolution of 4 km. Black triangles correspond to stations where surface-tethered drifting sediment traps were deployed (170 m, 270 m, 1000 m). Grey dots correspond to stations where Marine Snow Catcher (MSC) casts were performed at three depths (see Methods), and white dots to MSC casts performed at one depth (200 m). Black circles correspond to stations where the bottlenet profiles were performed between 2000 m and 200 m. b. Abundances of the five *nifH* phylotypes (Log10 *nifH* gene copies L^-1^) studied during the cruise over the transect (averages over the photic layer, ∼0-130 m).

## Results and discussion

### Surface conditions

The study region was characterized by typical oligotrophic waters (chlorophyll concentrations <0.15 µg L^-1^, DCM 100-180 m) east and west of Tonga, corresponding to the first group of stations S01, S02, S07 and S08 (Fig. 1a). The second group of stations (S03, S04, S05, S10, S11) was located in mesotrophic waters (chlorophyll >0.15 µg L^-1^, DCM 70-90 m) in the vicinity of the Tonga volcanic arc (Fig. 1a). Surface (0-50 m) nitrate concentrations were consistently close or below the detection limit (0.05 µmol L^-1^) throughout the transect, while phosphate concentrations were typically 0.1 µmol L^-1^ at oligotrophic stations and depleted down to detection limit (0.05 µmol L^-1^) at mesotrophic stations, likely due to consumption by higher plankton stocks (Fig. S1). Seawater temperature ranged from 23.1 to 27.3 °C in the mixed layer (0-15 m to 0-60 m) determined according to de Boyer Montégut et al. (2004)^27^ (Fig. S1).

The abundances of key diazotroph groups indicated that UCYN-A1 were present at high abundances throughout the photic layer (0-130 m) of the transect (average 3.2 x 10^8^ *nifH* gene copies L^-1^) (Fig. 1b), peaking at oligotrophic stations (average 1.1 x 10^9^ *nifH* gene copies L^-1^). *Trichodesmium* was the second most abundant group, particularly in the vicinity of Tonga (average 1.3 x 10^7^ *nifH* gene copies L^-1^), followed by UCYN-B (5.7 x 10^5^ *nifH* gene copies L^-1^). UCYN-C and Gamma-A (a non-cyanobacterial diazotroph) were also detected throughout the transect albeit at lower abundances (10^2^ to 10^4^ *nifH* gene copies L^-1^).

### How efficient are different diazotrophs groups to export?

We first examined the *nifH* gene community composition of particles collected in drifting sediment traps located at 170 m, 270 m and 1000 m at stations S05M and S10M (Fig. 1a). On average, 32% of the retrieved Amplicon Sequence Variants (ASV) corresponded to cyanobacteria genera over both stations (Fig. 2). The most abundant genus was *Trichodesmium* sp., but ASVs related to *Crocosphaera watsonii* (UCYN-B), *Candidatus* Atelocyanobacterium thalassa (UCYN-A) and *Katagnymene* spp. were also identified (Fig. 2). Genera related to DDAs (*Richelia* spp., *Calothrix* spp.) contributed little to the percentage reads. Besides autotrophic diazotrophs, 68% of ASVs were affiliated to non-cyanobacterial diazotrophs including Alpha-, Beta-, and Gammaproteobacteria at both stations.

**Figure 2.**
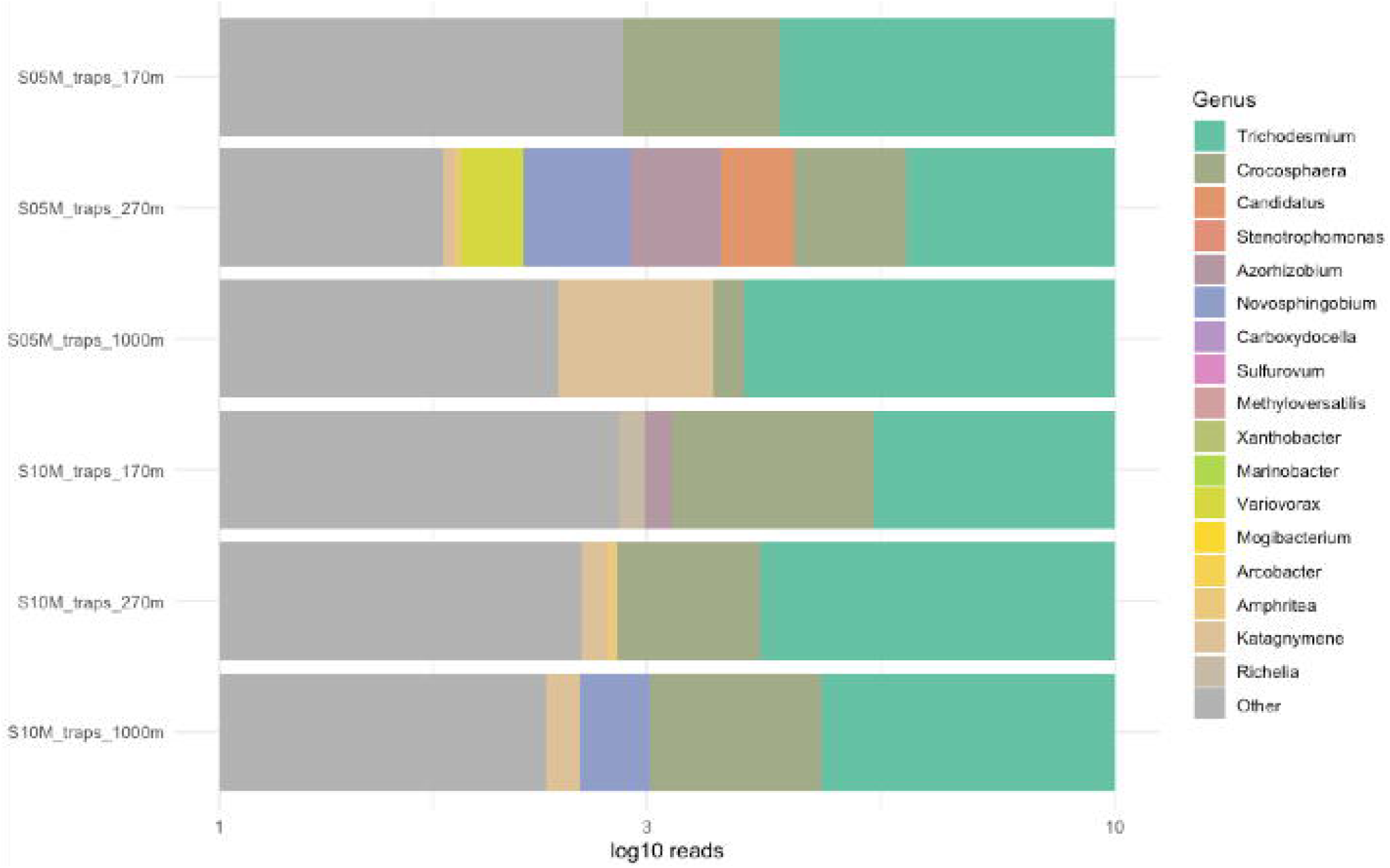
Relative abundance of diazotroph genera on sediment trap samples. The dendrogram corresponds to the clustering of samples generated by Bray-Curtis similarity. The legend indicates the *nifH* genus affiliation.

Amplicon sequencing provides relative abundances, which are not necessarily indicative of the abundance of the target microorganism in the natural environment^28,29^. Hence, to assess the export capacity of individual diazotroph groups, we quantified the abundance of five groups spanning different forms, sizes, lifestyle (symbiotic or not), using qPCR assays (UCYN-A1 symbiosis, UCYN-B, UCYN-C, *Trichodesmium* and Gamma-A; see Methods). DDAs were not quantified as they were almost not detected on microscopy inspections, pointing to their rarity in those waters at the time of the cruise. All five groups were detected at abundances ranging from 10^3^ to 10^8^ *nifH* gene copies L^-1^ of traps material, the highest being reported for UCYN-A1 and the lowest for UCYN-C (Fig. 3a,b). The diazotrophs assemblage exported to sediment traps generally reflected that of the photic layer, although some diazotrophs seemed to sink more efficiently than others: the highest diazotroph export fluxes were measured for UCYN-A1 at both sites and all depths of traps deployment (5.0 ± 1.1 × 10^8^ to 8.5 ± 2.0 × 10^9^ *nifH* gene copies m^-2^d^-1^), followed either by UCYN-B (2.2 ± 0.9 × 10^7^ to 7.5 ± 1.4 × 10^7^ *nifH* gene copies m^-2^d^-1^) or *Trichodesmium* (2.5 ± 1.9 × 10^6^ to 1.2 ± 0.3 × 10^8^ *nifH* gene copies m^-2^) depending on station and depth (Fig. 3a,b). Gamma-A and UCYN-C were also exported, albeit at lower rates (∼10^5^ and ∼10^6^ *nifH* gene copies m^-2^d^-1^, respectively).

**Figure 3.**
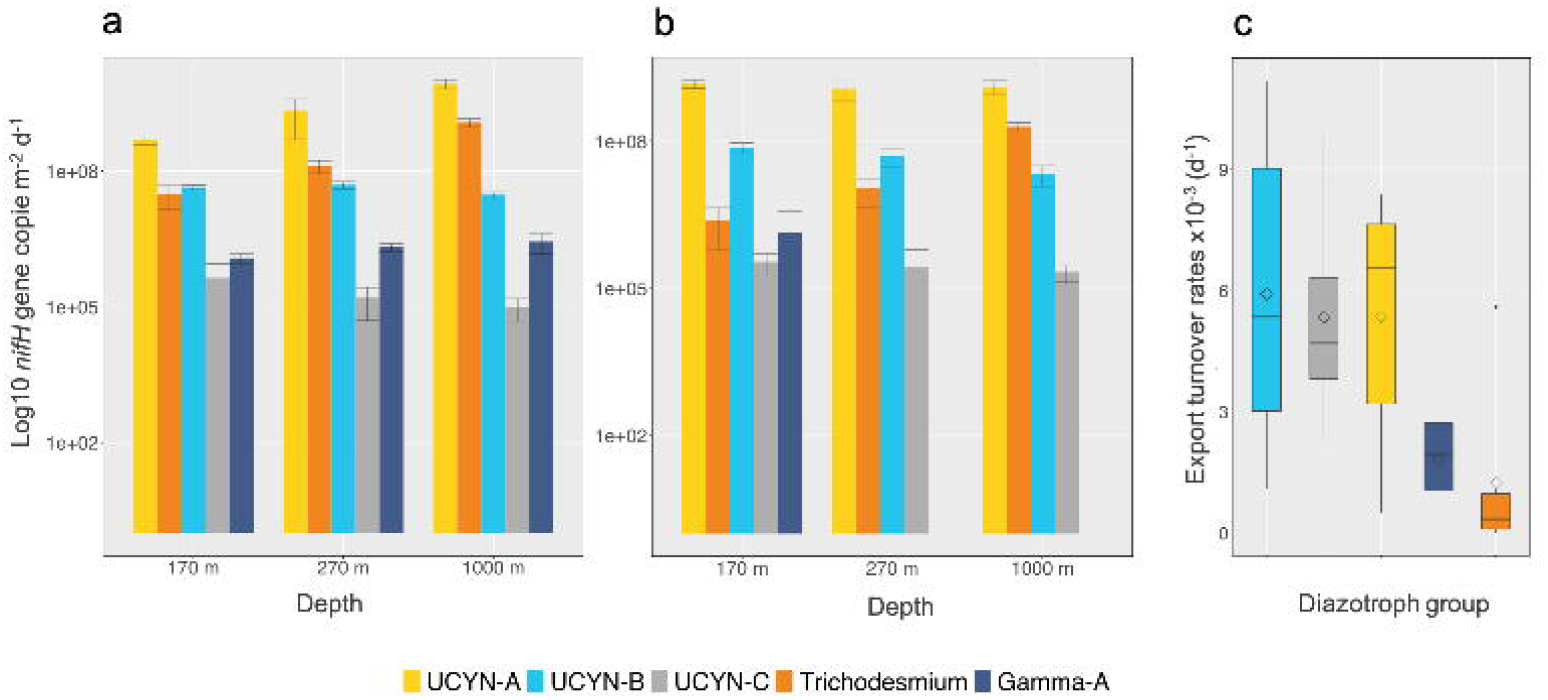
a. Export flux (*nifH* gene copies m^-2^ d^-1^) of the five diazotroph groups targeted by qPCR (UCYN-A1 symbiosis, UCYN-B, UCYN-C, *Trichodesmium* and Gamma-A) in sediment trap samples at 170 m, 270 m and 1000 m at stations S05M and b. at S10M. Error bars represent strandard deviations from triplicate aliquote analyzed in duplicates. c. Export turnover rates (d^-1^) of the same diazotroph groups (average of the three depths and the two stations for each group).

Specific export turnover rates provide information on the rate at which diazotrophs are “lost” from the photic layer due to export (Turnover rate (d^-1^) = export flux/abundance in the photic layer). The range of calculated rates (0.7 ± 1.2 x10^−5^ to 11.2 ± 1.9 x 10^−3^ d^-1^) spans through two orders of magnitude and varied depending on the diazotroph group, depth and station considered. On average (over all stations and depths), the export turnover rate of UCYN (5.5 ± 3.1 x 10^−3^ d^-1^) was ca. four times higher than that of *Trichodesmium* (1.3 ± 2.2 x 10^−3^ d^-1^) (Fig. 3c), pointing to preferential export of UCYN groups relative to the filamentous *Trichodesmium*. Among UCYN groups, the highest export turnover rate was measured for UCYN-B, followed by UCYN-C and UCYN-A1, although differences were not significant between groups (Mann-Whitney test, p<0.05). The export turnover rate of Gamma-A (average 1.8 ± 1.5 x 10^−3^ d^-1^) was intermediate between that of *Trichodesmium* and UCYN groups.

Epifluorescence microscopy confirmed the presence of phycoerythrin-containing UCYN-B and UCYN-C-like cells in sediment trap samples (Fig. 4a-f). Scanning electron microscopy revealed that they were recurrently found embedded in large organic aggregates, or organized into clusters of tens to hundreds of cells linked by an extracellular matrix (Fig. 4g-l), which was further confirmed by Alcian blue straining (Fig. S2). *Trichodesmium* was also observed in all samples, mostly as free filaments, but intact colonies were observed in sediment trap samples especially at 1000 m at both stations. Gamma-A and UCYN-A1 cannot be visualized by these techniques and are thus assessed solely on the basis of qPCR counts (above).

**Figure 4.**
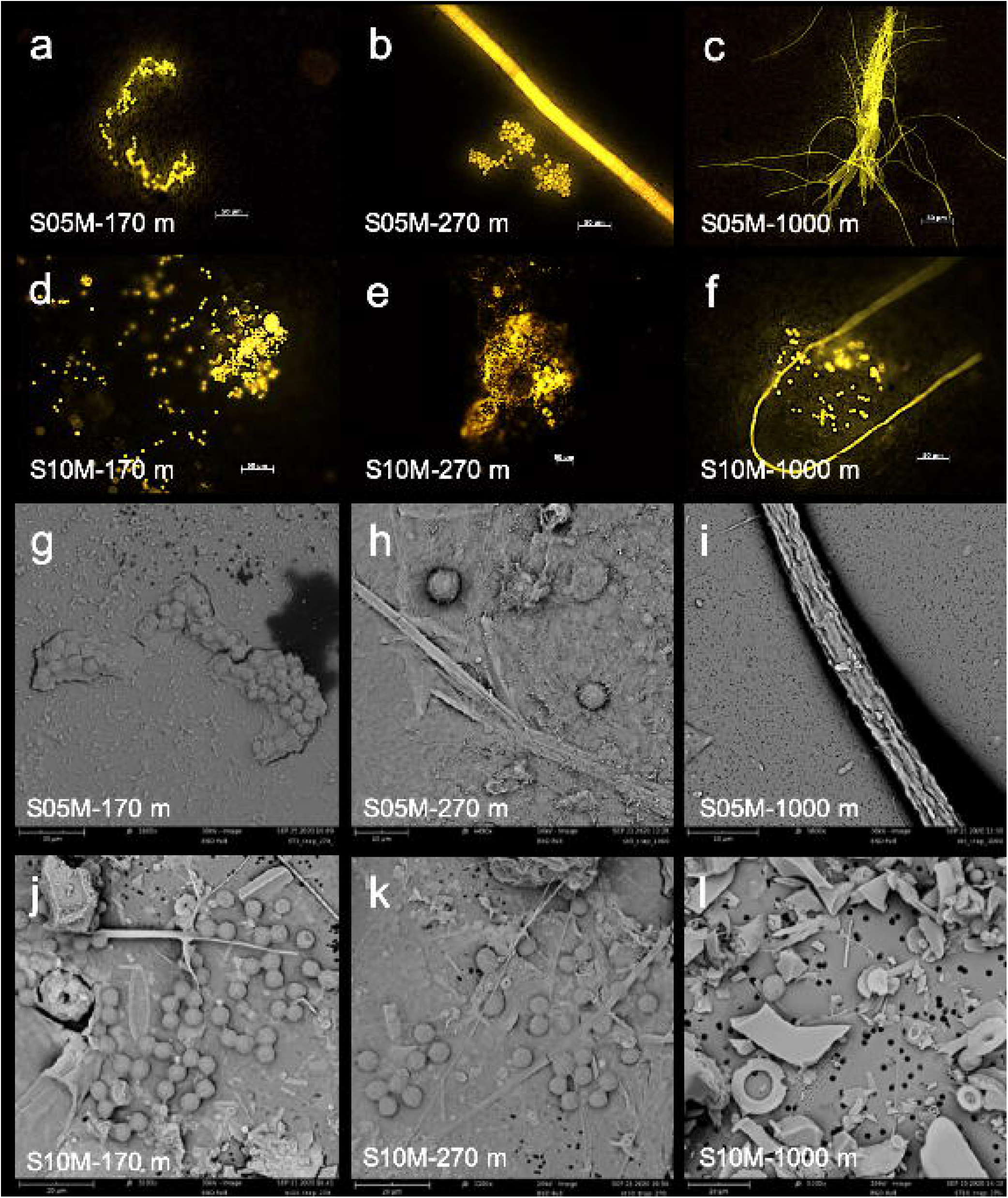
Microscopy images showing examples of phycoerythrin-containing UCYN-like cells and *Trichodesmium* in sediment trap samples collected at 170 m, 270 m, and 1000 m at stations S05M and S10M. a-f. Images taken by epifluorescence microscopy (green excitation 510–560 nm, scale bar: 50 µm). k-l. Image taken by scanning electron microscopy (SEM).

Pigment concentrations measured in sediment trap samples at both stations indicate that total pigment concentrations and diversity decreased with depth. Apart from the degradation pigments phaeophorbide and phaeophytin, zeaxanthin, a biomarker of cyanobacteria predominated at all depths (Fig. S3). The Chlorophyll *a* : phaeopigments ratios were elevated (average 1.8, range 0.8-3.), indicative of fresh organic matter in sediment trap samples.

### Contribution of diazotrophs to PON export fluxes

Sediment trap-based PON export fluxes at 170 m, 270 m and 1000 m ranged between 1.6 ± 0.2 and 6.1 ± 1.2 mg N m^-2^ d^-1^ at S05M and 5.2 ± 0.1 and 13.4 ± 1.3 mg N m^-2^ d^-1^ at S10M (Table 1). Fluxes attenuated with depth, with transfer efficiencies of 0.02 and 0.03 between 170 m and 1000 m at both stations. The N export efficiency (Ne-ratio, the amount of PON export at the base of the euphotic zone over the integrated N_2_ fixation rate) was 18% and 53% at stations S05M and S10M. The C:N ratios in exported material was lower (5.2) than that of the surface (7.5, integrated value over the photic layer) at station S10M, pointing to a preferential export of PON relative to POC.

**Table 1.** Particulate nitrogen (PON) export fluxes at stations S05M and S10M at 170 m, 270 m and 1000 m, and contribution of diazotrophs to total PON export.

| Station/Depth (m) |  | 170 m | 270 m | 1000 m |
| --- | --- | --- | --- | --- |
| PON export flux (mg N m<sup>-2</sup> d<sup>-1</sup>) |  | 4.9±0.6 | 6.1±1.2 | 1.6±0.2 |
| Diazotroph-PON (mg N m<sup>-2</sup> d<sup>-1</sup>) | S05M | 0.9 | 2.3 | 1.6 |
| Contribution diazotrophs (%) |  | 18 | 38 | 100 |
| PON export flux (mg N m<sup>-2</sup> d<sup>-1</sup>) |  | 13.4±1.3 | 9.7±0.6 | 5.2±0.1 |
| Diazotroph-PON (mg N m<sup>-2</sup> d<sup>-1</sup>) | S10M | 1.2 | 1.2 | 2.7 |
| Contribution diazotrophs (%) |  | 9 | 12 | 52 |

The proportion of PON attributed to diazotrophs and collected by sediment traps was evaluated based on qPCR results and estimated PON content per diazotroph (see Methods). The PON flux attributed to diazotrophs ranged from 0.9-2.3 mg N m^-2^ d^-1^ at S05M and from 1.2-2.7 mg N m^-2^ d^-1^ at S10M, accounting for 18% (170 m) to 100% (1000 m) and 9% (170 m) to 52% (1000 m) of the total PON export flux at stations S05M and S10M, respectively (Table 1). *Trichodesmium* was the major contributor of the PON attributed to diazotrophs at 1000 m at both stations, whereas UCYN dominated at shallower depths (Table S1). Despite non-cyanobacterial diazotrophs were present in exported material (68% of sequences), their C and N composition remains unknown due to the poor availability of cultured representatives. Thus, quantifying their contribution to export fluxes is not possible at this stage. In addition, sequencing data also revealed the presence of *Katagnymene* spp., which are large (>100-1000 µm) and likely have high C and N content. Hence the contribution of diazotrophs to exported PON reported here likely represent a lower-end estimate.

### The fate of diazotrophs in the mesopelagic ocean

We further assessed the fate of the exported diazotroph community by collecting fresh particles in mesopelagic waters using the Marine Snow Catcher (MSC, see Methods). In essence, the MSC allows separating non-sinking particles from slow sinking and fast sinking particles^30^ (see Methods). These different fractions can hence be studied individually. MSC samples were collected at the same depths as those of sediment traps (170, 270, 1000 m) at stations S05M and S10M and in addition at seven other stations (Fig. 1a).

The diazotroph community composition (based on *nifH* amplicon sequencing) in the various fractions generally mirrored that of traps at stations S05M and S10M with 32-54% of sequences affiliated with cyanobacteria and a prevalence of *Trichodesmium* (Fig. 5). This was also the case at stations S03, S04, S07 and S11, but not at stations S01, S02 and S08, where sequences assigned to *Candidatus Atelocyanobacterium thalassa* and *Crocosphaera watsonii* were generally more abundant than *Trichodesmium* sequences. As in sediment traps, a large number of *nifH* sequences (46-58%) were affiliated to non-cyanobacterial diazotrophs, whose composition was overall consistent with that of the traps, although some classes not detected in sediment traps (Epsilonproteobacteria and Clostridia) were present in all MSC fractions. In general, the total number of *nifH* gene reads was consistently 10-25% higher in the slow and fast sinking fractions compared to the suspended fraction at all stations. This trend was mainly driven by non-cyanobacterial sequences that are suspected to be attached to particles^31^, and to a lesser extent by sequences affiliated with cyanobacteria.

**Figure 5.**
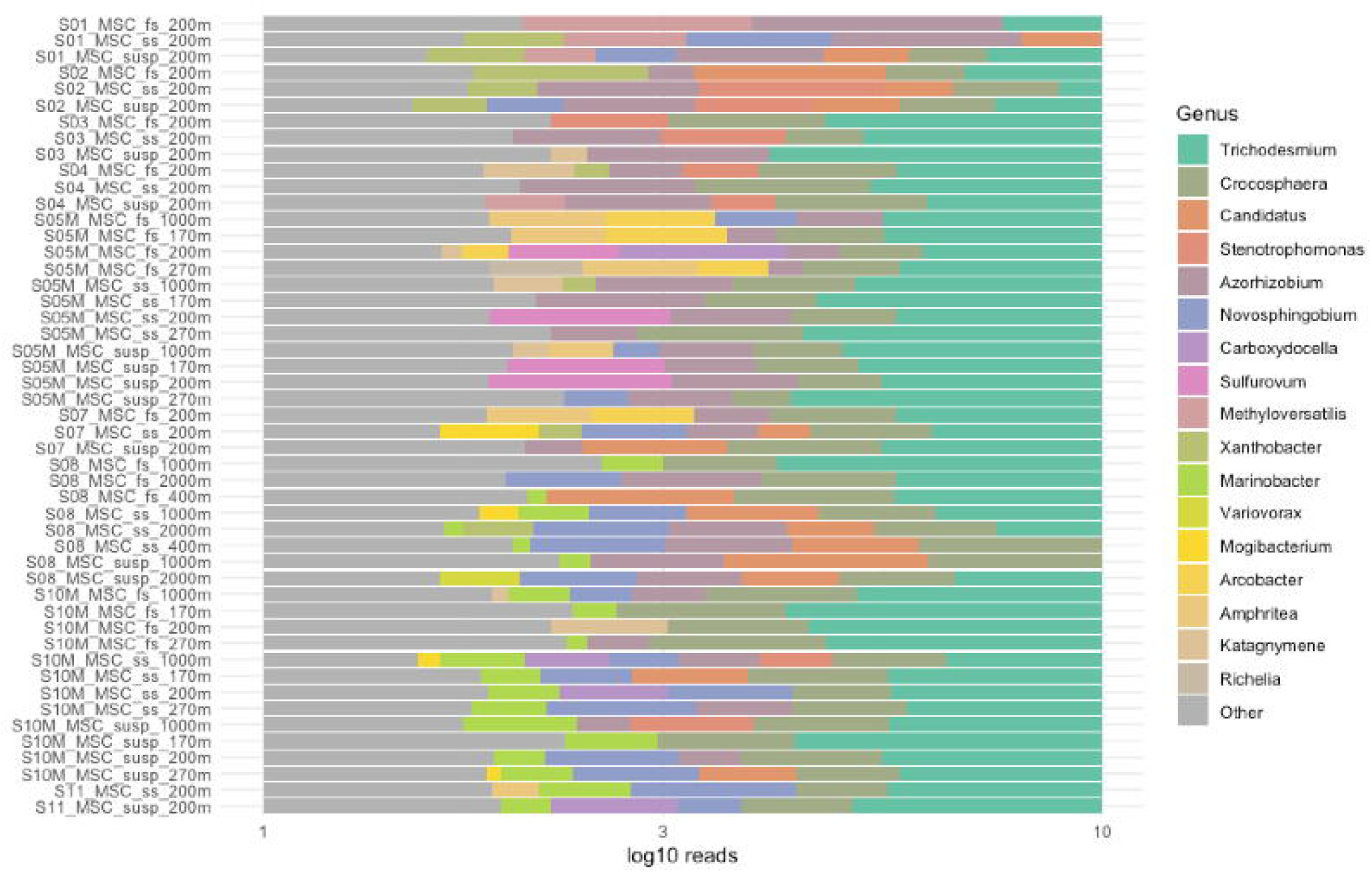
Relative abundance of diazotroph genera from marine snow catcher samples. The dendrogram corresponds to the clustering of samples generated by Bray-Curtis similarity. The legend indicates the *nifH* genus affiliation.

As in sediment traps, five diazotroph groups were quantified in the suspended, slow and fast sinking fractions to more finely assess their sinking dynamics over short time scales (h). Overall, UCYN-A1 and *Trichodesmium* were the most abundant groups in the mesopelagic MSC samples (170 m to 1000 m), and their numbers varied in parallel with the abundance of their population in the photic layer (Fig. 6). UCYN-A1 were the most abundant at oligotrophic stations (S01, S02, S07, S08, chlorophyll <0.15 µg L^-1^), while *Trichodesmium* peaked at mesotrophic stations (Chlorophyll a >0.15 µg L^-1^, Fig. 1). Conversely, the abundance of UCYN-B, UCYN-C and Gamma-A in mesopelagic waters was poorly correlated with their photic layer abundance.

**Figure 6.**
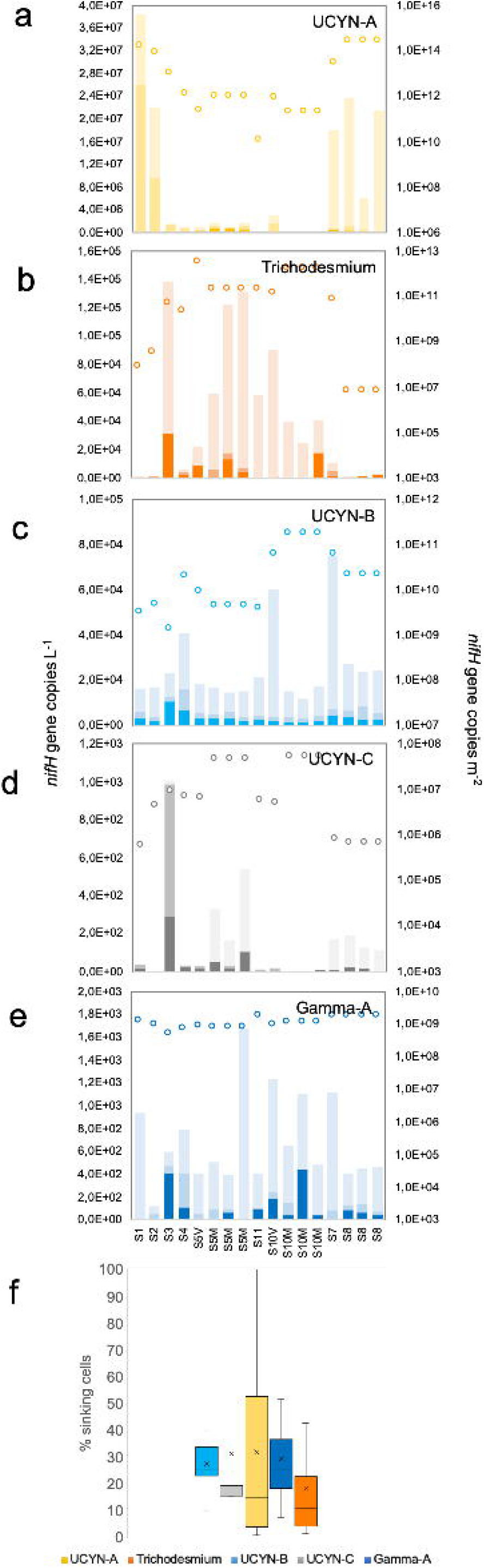
a-e. Quantification of the five diazotroph groups targeted by qPCR in Marine Snw catcher samples: UCYN-A1 symbiosis: *Trichodesmium*, UCYN-B, UCYN-C, and Gamma-A in the suspended (dark color), slow sinking (medium color) and fast sinking (light color) pools of the MSC samples across the transect (left Y axis, *nifH* gene copies L^-1^). Dots represent the integrated pool of each phylotype (right Y axix, *nifH* gene copies m^-2^) in the photic (∼0-130 m) layer. f. Percentage of sinking versus non-sinking diazotrophs for each group.

The MSC suspended fraction accounted for the most abundant pool of diazotrophs at all stations, followed by either the slow or the fast sinking pools, depending on groups (Fig. 6a-e). At the mooring stations S05M and S10M, the total concentration of diazotrophs and the diazotroph community composition remained generally constant at the three sampled depths (170 m, 270 m and 1000 m), consistent with the results observed in sediment traps. To evaluate the export potential of each individual diazotroph group, we calculated the ratio of sinking versus non-sinking cells (the sum of the slow and fast sinking fractions over the total). The proportion of sinking cells varied widely across the transect, but was the highest for UCYN-A1 (32 ± 33 %), UCYN-C (31 ± 38 %), Gamma-A (29 ± 19 %) and UCYN-B (27 ± 11 %), and the lowest for *Trichodesmium* (18 ± 20 %) (Fig. 6f). Among the sinking fractions, UCYN-B and Gamma-A were generally equally distributed among the slow and fast sinking fractions, whereas the majority of *nifH* gene copies were found in the fast sinking fraction for UCYN-A1 (59%), UCYN-C (62%) and *Trichodesmium* (67%). Taken together, these results indicate that over short time scales (2 h, the conventional settling of particles following MSC deployment^30^): i) small UCYN (few µm) generally sink more efficiently than large *Trichodesmium* (>100 µm), ii) when sinking, *Trichodesmium* sinks fast, iii) UCYN-A1 and UCYN-C sink faster than UCYN-B and Gamma-A. A detailed imaging study performed on 170 m MSC samples at the mooring stations S05M and S10M indicates that the 50-80% of phycoerythrin-containing UCYN in the fast sinking fraction were organized into aggregates of tens to >250 cells measuring 30 to >100 µm, while the majority (60-95%) of UCYN were free living in the suspended fraction (Figure S4). This indicates that under minimum turbulent agitation, UCYN quickly (within 2 h) form aggregates large/dense enough to sink.

Concentrations of PON averaged across all stations 1.85 ± 0.49 µg L^-1^, 0.25 ± 0.13 µg L^-1^ and 0.26 ± 0.08 µg L^-1^ within the suspended, slow sinking and fast sinking fractions (Fig. S5a). PON in the fast sinking fraction contributed 11 ± 2 %, while the slow sinking and suspended fractions contributed 11 ± 4 % and 79 ± 5 % in terms of total PON. Thus, ∼22% of the PON was sinking out of the upper part of the MSC within 2h during our study. We converted transparent expolymeric particles into C (TEP-C), that revealed higher concentrations in the suspended fraction compared to the fast sinking fraction at all stations (TEP-C concentrations were null in the SS fraction) (Fig. S5b). The TEP-C:POC ratio was generally higher in the suspended fraction compared to the fast sinking. This indicates that, despite UCYN were embedded in TEPs in FS samples, cells rather than TEPs were the major contributor of the POC pool in the FS fraction.

Finally, we quantified diazotrophs on vertical profiles spanning the water column between 200 m and 2000 m by using a Bottle-net mounted on the CTD rosette frame at six stations across the transect (Fig. 1a). The bottle-net consists of a 20-μm conical plankton net housed in a cylindrical PVC pipe^19^. The top cover is opened at the desired bottom depth (2000 m) of the tow, remains opened during the ascent of the rosette, and closed again at the upper depth (200 m) of the water column to be sampled. This results in one integrated sample of 200 m to 2000 m per deployment. Overall, bottle-net tows confirm that diazotrophs are consistently present in this deep ocean layer, with concentrations averaging 1.4 x 10^7^ *nifH* gene copies m^-2^. Among the groups targeted by qPCR, the community was primarily dominated by UCYN-A1 (63% on average over all sampled stations) and *Trichodesmium* (27%), and secondarily by UCYN-B (9%) (Fig. 7), generally mirroring the diazotroph community in surface water (these numbers are conservative and may be underestimated as some individual UCYN pass through the 20 µm mesh net of the bottle-net). The 2000-200 m stock was relatively constant among stations, except at the most oligotrophic station S08, where it was lower by two orders of magnitude than that of other stations.

**Figure 7.**
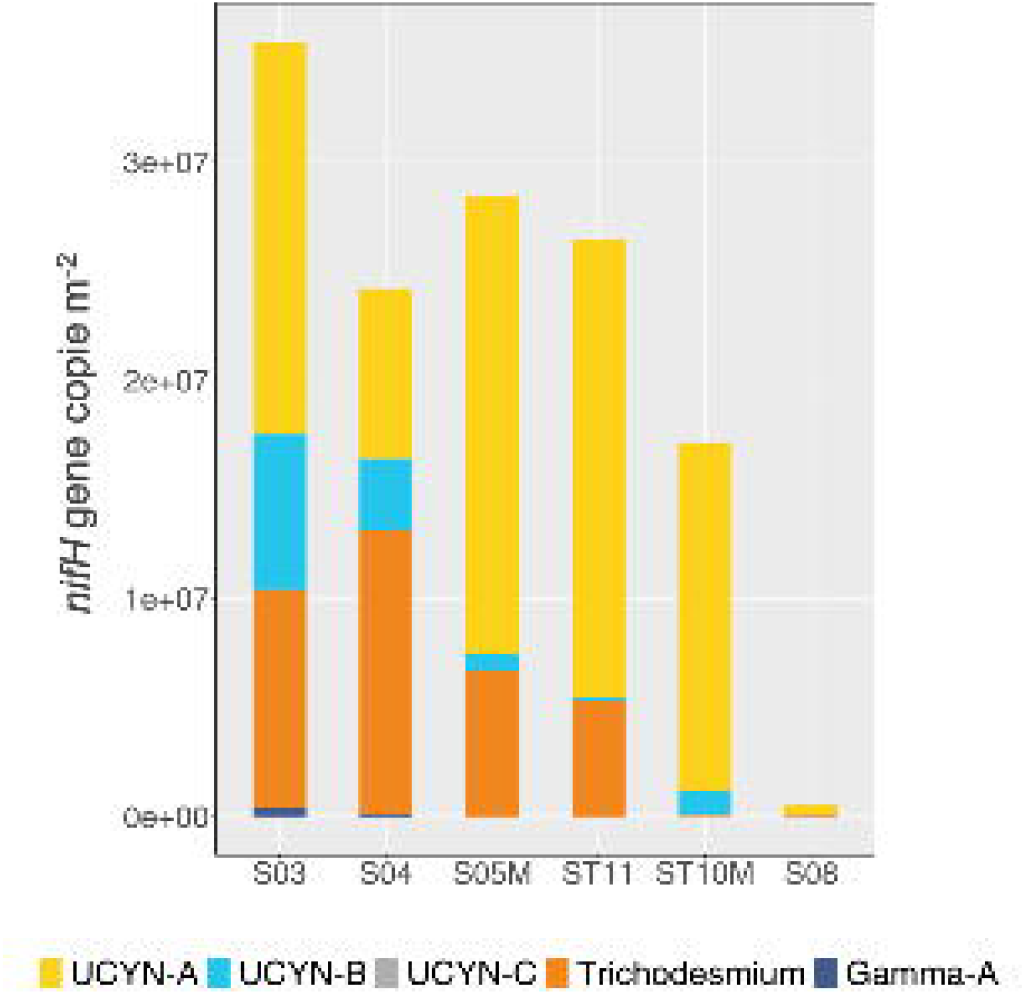
Abundance (*nifH* gene copies m^-2^) of the five diazotroph groups targeted by qPCR (UCYN-A1 symbiosis, *Trichodesmium*, UCYN-B, UCYN-C, and Gamma-A) in the bottle-net samples across the transect, representing an integrated sample from 200 m to 2000 m.

### Beyond subtropical south Pacific waters

To assess whether the sinking of globally-distributed diazotrophs down to mesopelagic waters is a widespread feature, we explored the presence of diazotrophic cyanobacterial genomes using *Tara* Oceans metagenomes collected from other ocean basins^32,33^. Among all the stations for which metagenomic samples from both surface and mesopelagic were available, we selected those ones showing significant abundances of *nifH* metagenomic reads in surface waters (5 m)^32^. This led to a total of five stations located in the South Atlantic and North and South Pacific oceans where we also explored the presence of diazotrophs, at the genome level, in the mesopelagic (300 to 800 m) (Fig. 8). Recruitment of metagenomic reads against representative genomes of the diazotrophic community (*Trichodesmium, Richelia*, UCYN-B and UCYN-A1 and UCYN-A2; Table S2) across different size-fractions in surface and mesopelagic waters show that diazotrophs in general were systematically detected in mesopelagic waters at all five stations (Fig. 8; Table S3 ; see Methods). As in the WTSP, every specific group was detected at depth when present in surface, except when abundances in surface were very low (<6.7 reads/total 100,000 reads). Accordingly with their cell size, *Trichodesmium, Richelia*, UCYN-B and UCYN-A2 reads were recovered in the size fraction >3 µm at mesopelagic depths, whereas the UCYN-A1 symbiosis was recovered in the 0.2-3 µm and 0.8-3 µm size fractions. Overall, this metagenomic analysis shows that the export of diazotrophs through gravitational settling is not restricted to the WTSP and is a widespread phenomenon in the tropical ocean.

**Figure 8.**
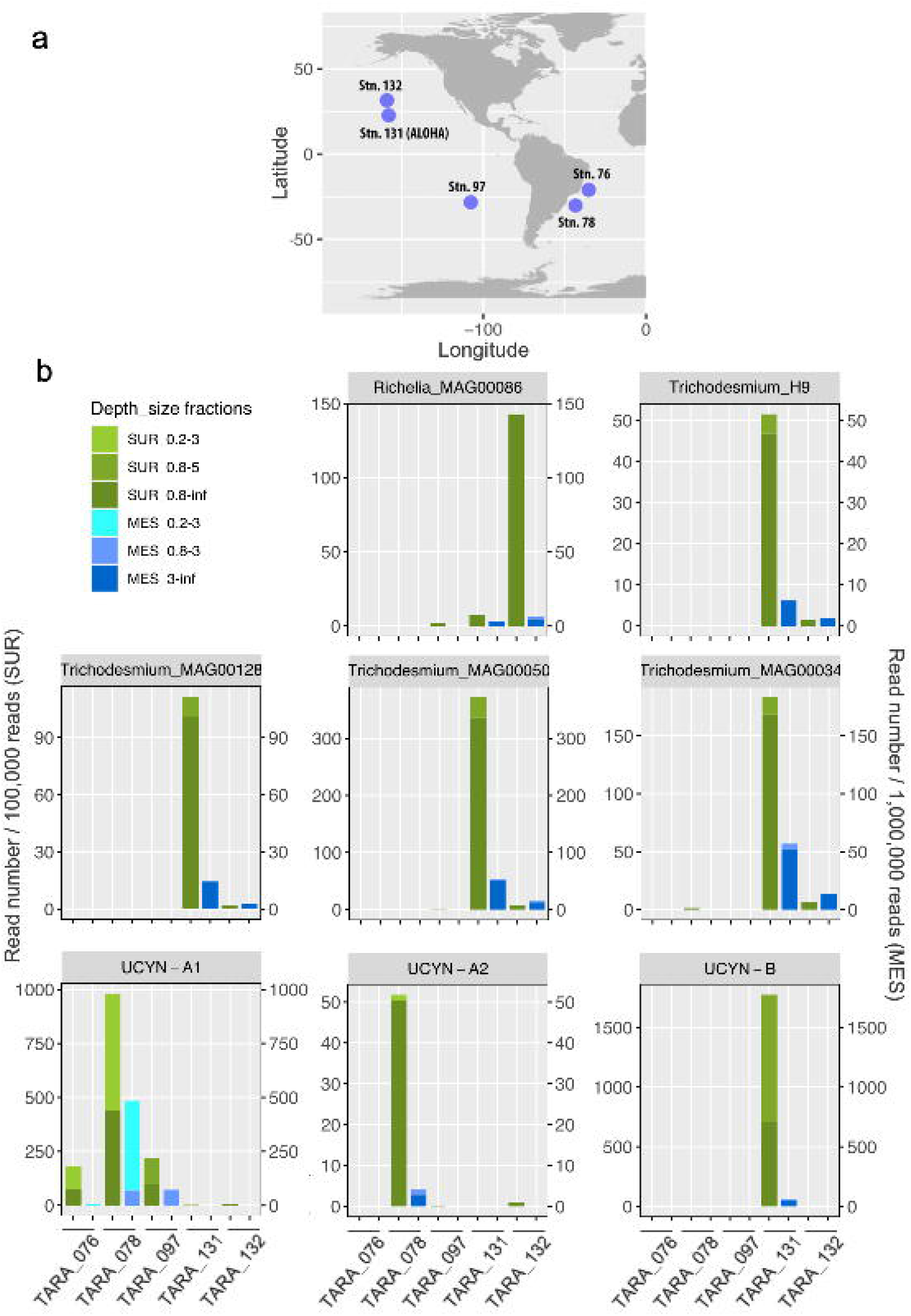
Distribution of diazotrophic cyanobacteria in surface and mesopelagic waters during the *Tara* Oceans expedition. A. Geographical location of the *Tara* oceans stations used in this study. Only stations in which diazotrophic cyanobacteria were present in surface and metagenomes were available both from surface and mesopelagic waters were selected. B. Abundance of metagenomic reads recruited against different diazotrophic cyanobacteria genomes in surface and mesopelagic samples. See Table S2 for the complete dataset. Note that to better visualize the data, read abundance was expressed as number of reads per 100,000 total reads for surface samples (left axis) and per 1,000,000 total reads for mesopelagic samples (right axis).

### Biogeochemical implications

Excluding silicified DDAs^11^, diazotrophs have seldom been regarded as important contributors to organic matter export. Yet, our results provide clear evidence that they are present and ubiquitous in the mesopelagic ocean and they contribute significantly to the PON export fluxes in the WTSP. They present, however different behaviors.

We showed that diazotroph cell sizes is not necessarily a key variable controlling the diazotrophs’ ability to sink out the euphotic zone. Small UCYN (1-8 µm) displayed the highest export turn-over rates ∼10^−3^ d^-1^, in the range or higher than those reported for ballasted phytoplankton groups such as diatoms and coccolithophores^34^. Within UCYN, UCYN-A1 clearly dominated the diazotroph community targeted by qPCR in mesopelagic waters of the WTSP. Although the UCYN-A1 symbiosis has been detected once in mesopelagic waters of the South Atlantic Ocean^32,35^, their capacity to leave the photic layer and sink throughout the mesopelagic zone down to 1000 m (Fig. 3, 8) has not been thoroughly explored before. As UCYN-A rank among the most abundant diazotrophs in the global ocean^12^ and span from tropical to polar waters^36,37^, we suggest that they potentially play a significant contribution to ocean organic matter sequestration. Some UCYN-A ecotypes (e.g. UCYN-A2) live in obligate symbiosis with a calcified coccolithophore^38^, hence their export is likely enhanced by this ballast mineral increasing the density of sinking particles to which they are associated. In future studies, better constrains on UCYN-A sedimentation rates and aggregation processes would be of primary importance to assess their role in particle fluxes and cycling.

UCYN-B and -C are free-living unicellular cyanobacteria and their export pathway therefore does not rely on mineral ballasted hosts. Our results indicate that the sinking of such small cells is made possible through the aggregation of UCYN into large-sized aggregates (30->100 µm) of tens to hundreds of cells, large/dense enough to sink, in accordance with a previous mesocosm study^21^. The presence of UCYN-B in sediment traps is also in accordance with previous reports of *Crocospheara watsonii* sequences at station ALOHA either just below the euphotic zone^22^, or as deep as 4000 m^39^. The visualization of our sediment trap samples shows that they can be subsequently embedded in mixed aggregates with other cells and debris (Fig. 4, S2). The UCYN-B large (>4 µm) ecotype, which was dominating in this study (data not shown), produces large amounts of C-rich transparent exopolymeric particles (TEP) at rates one to two order of magnitude higher than that of diatoms and coccolithophorids^40^, probably in response to nutrient limitation and excess light^41^. In cultures of *Crocosphaera watsonii*, TEP account for ∼22% of the particulate C pool^40^. Hence, TEP produced by UCYN-B do not only provide a matrix for the formation of large aggregates, but may also account for a significant fraction of C export. TEP are indeed greatly enriched in C relative to N (C:N ∼25^40^).

*Trichodesmium* was generally the second contributor to the diazotroph community targeted by qPCR in mesopelagic waters after UCYN-A1, which contradicts the common assertion that they are entirely remineralized in the euphotic layer^4,13^. Several potential mechanisms could explain the presence of *Trichodesmium* in mesopelagic waters. *Trichodesmium* colonies can migrate vertically to exploit the deep phosphate stock (Villareal and Carpenter 2003) (the phosphacline was around 200 m in our study region at the time of the cruise). According to this theory, they overcome their positive buoyancy by fixing C that result in carbohydrate ballasting. However, Walsby (1978)^42^ observed that 100% of the gas vacuoles of *Trichodesmium erythraeum* (the most abundant species in this study) collapse at depths between 105 m and 120 m, resulting in a loss of their buoyancy. Hence, *Trichodesmium* could be locked in a persistent and irreversible downward trajectory. Alternatively, Berman-Frank et al.^43,44^ have shown that programmed cell death induces internal cellular degradation, gas-vacuole loss and increased production of TEP, resulting in an increase in the vertical flux of *Trichodesmium*^45^. Whatever the mechanism behind the vertical flux of *Trichodesmium*, we report here higher abundances at 1000 m compared to those at shallower depths at both stations, consistent with microscopic observations showing intact colonies at 1000 m. This likely results from a spatio-temporal decoupling between production and export^46^. Despite sinking less efficiently than UCYN in the aggregates, they are much larger and contain more C and N per filament than UCYN^12^, and accounted for a significant fraction of the diazotroph-attributed PON export in this study, especially at 1000 m (Table S1), which is in accordance with previous studies reporting intact *Trichodesmium* colonies down to 3000-4000m^16,19^.

More than 90% of the organic matter sinking below the euphotic zone is respired before it reaches a depth of 1000 m^7^. Fast sinking particles will therefore theoretically make a greater contribution to the deep ocean flux than slow sinking particles, since the latter will be rapidly recycled at shallow depths. The relative similarity of the taxonomic diazotroph community composition in mesopelagic waters sampled by three independent sampling approaches (traps, bottle-net, and MSC) compared to the diazotrophic composition of the euphotic layer suggests a rapid export mode of diazotrophs. This is further confirmed by i) the high proportion of diazotroph groups quantified in the MSC fast sinking fraction, ii) the whole cells and colonies and high Chlorophyll *a* : pheopigments ratios in sediment traps, indicative of healthy phytoplankton (Fig. 4; S2). Finally, iii) while PON export fluxes were attenuated with depth in our study, which is a classical feature in the oligotrophic ocean^47^, the diazotroph export fluxes were not, suggesting a high transfer efficiency to the deep ocean, and thus sinking velocities high enough to escape short-term remineralization. Indeed, a parallel study performed during the same cruise^48^ reveals significant N_2_ fixation rates in sediment traps and MSC samples, suggesting that part of the diazotroph community in mesopelagic waters sunk fast enough to remain alive at mesopelagic depths. Bar-Zeev et al.^45^ reported sinking velocities of *Trichodesmium erythraeum* of ∼200 m d^-1^. More recently, Ababou et al.^49^ reported sinking velocities of *Trichodesmium erythraeum* and UCYN aggregates formed in roller tanks of 92 ± 37 m d^-1^ and 333 ± 176 m d^-1^, respectively. We calculated that *Trichodesmium* sinking at those rates (92-200 m d^-1^) would take 5 to 10 days to reach 1000 m, whereas UCYN would take 3 days, which would be compatible with finding active cells observed at those depths^48^.

## Conclusion

Our findings challenge the common assumption that the fate of diazotroph-derived production is constrained to the surface layer. They provide a significant advance in defining the role of diazotrophs on an important planetary carbon flux, the biological carbon pump, and bring new insights into the species-specific export of diazotrophs in the ocean, revealing the previously-unseen role of UCYN and *Trichodesmium* into the overall export fluxes. Diazotrophs being not restricted to the tropical ocean, it would be interesting in future studies to explore the direct gravitational export of diazotrophs in temperate and polar waters.

Direct export through gravitational settling of diazotrophs is most likely only the tip of the iceberg and diazotrophs can potentially be exported through secondary pathways: diazotrophs release in seawater 10–50% of recently fixed N_2_ (referred to as Diazotroph Derived, DDN) as NH_4_^+^ and dissolved organic N (DON)^50,51,52^. This DDN is potentially available for assimilation by the surrounding phytoplanktonic communities, supporting their growth and leading a potential secondary (indirect) export pathway of diazotroph-derived OC^6,21,53^. Moreover, DDN is also transferred to zooplankton, which produces dejections termed fecal pellets, that sink fast and play a major role in OC export to the deep ocean^54^. In future studies, there is an urgent need to develop appropriate approaches to decipher diazotroph export pathways (direct *vs* indirect) if we are to understand the role of N_2_ fixation in the biological carbon pump. This is an especially pressing question given that current climate models predict an expansion of the oligotrophic gyres (60% of our oceans)^23,24^, where diazotrophs thrive. N_2_ fixation will thus likely be crucial to supporting primary productivity and export in the future ocean.

## Methods

Samples were collected during the TONGA cruise (Fig. 1, DOI: 10.17600/18000884) in the tropical South Pacific Ocean onboard the R/V *L’Atalante* from November 1 to December 5 2019. We collected suspended and sinking particles in the mesopelagic zone using three complementary devices: surface-tethered drifting sediment traps^55^, MSC and Bottle-net. Additionally, water column samples were collected from the euphotic layer using Go-Flo bottles mounted on a trace metal clean rosette to quantify the stocks of the major groups of diazotrophs in the photic layer.

### Satellite products and photic layer sampling

Satellite-derived surface chlorophyll *a* concentration during the TONGA cruise (November 2019) was accessed at http://oceancolor.gsfc.nasa.gov (MODIS Aqua, 4 km, 8-days composite, level 3 product). Vertical 0-150 m depth profiles were performed at each station using a trace metal clean titanium rosette of Go-Flo bottles equipped with a fluorometer and temperature, conductivity and oxygen sensors. Seawater samples were collected from 5 depths (75, 50, 20, 10, 1% surface irradiance levels) to quantify the stock of major groups of diazotrophs in the photic layer by quantitative PCR (qPCR) as described below. Additionally, primary production and N_2_ fixation rates were measured in triplicates at the same six depths at the locations of the traps deployments (S05M and S10M), as described in ^56, 57^. N_2_ fixation rates were measured at two occasions (on day 1 and day 3 of the traps deployment). The average value between both profiles was used to calculate the e-ratios (POC export:primary production) and the contribution of N_2_ fixation to primary production rates.

### Sediment trap deployment and sample analyses

A surface tethered mooring line (∼1000 m long) was deployed at stations S05M (21.157°S;175.153°W, 5 days) and S10M (19.423°S;175.133°W, 4 days). The line was equipped with sediment traps (KC Denmark®) at 3 depths: 170 m (corresponding to the base of the photic layer), 270 m and 1000 m. Each trap was composed of four particle interceptor tubes (PITs) mounted on a cross frame (collecting area of 0.0085 m^2^, aspect ratio of 6.7). Two of the tubes were used for this study: one tube was used for biogeochemical analyses (hereafter referred to as ‘Biogeo’ tube), and one for microbiological analyses (‘Microbio’ tube). Prior to deployment, the tubes were filled with 0.2 μm filtered seawater with added saline brine (50 g L^-1^). Borate-buffered formalin (5%) was also added to the Biogeo tube to prevent in situ microbial decomposition^55^. After recovery, the density gradient was visually verified, and the PITs were allowed to settle for 2 h before the supernatant seawater was carefully removed with a peristaltic pump. The remaining water containing the sinking material was transferred to a chlorohydric acid-washed container, while being screened with a 500 μm mesh to remove swimmers (zooplankton that actively entered the traps)^58^. Subsequently, samples were split into 12 aliquots. A triplicate set of aliquots were filtered onto 25-mm diameter Supor filters for *nifH* sequencing and *nifH* qPCR as described below. Another triplicate set of aliquots were filtered onto 25-mm diameter combusted (4h, 450°C) glass microfiber filters (Whatman GF/F), which were subsequently dried for 24 h at 60°C, pelleted and from which particulate N (PON) and C (POC) were analyzed by EA-IRMS (Elemental Analyzer-Isotope Ratio Mass Spectrometry) using an Integra 2 (Sercon) mass spectrometer. Lastly, a triplicate set of aliquots was filtered on GF/F filters for further pigment analyses^59^, and another triplicate filtered on 1 µm polycarbonate filters for microscopic analyses (see below for methods).

### MSC deployment

Suspended and sinking particles were sampled using a MSC at 3 depths at the mooring stations S05M and S10M (170 m, 270 m and 1000 m), and at 200 m, 400 m and 1000 m at S8. Additionally, the MSC was deployed at 200 m at S01, S02, S03, S04, S05V, S07, S10V and S11 (Fig. 1). The MSC is a large volume water sampler (100 L) that collects sinking particles with minimal turbulent agitation^60^. Upon recovery, the MSC is placed on deck for 2 h while any particles present settle onto the base of the bottom 7 L chamber. After each recovery, the MSC is conventionally left on deck for 2 h allowing particles to settle^30^. Fast sinking particles were thereby deposited collected in a dedicated plate at the bottom of the MSC, slow sinking particles were collected from the 7-litre compartment above the plate, and the non-sinking (or suspended) fraction was collected from the upper part of the MSC^30^. All three fractions were collected separately (4 L for the suspended, 1.2 L for the slow sinking). For the fast sinking fraction, the entire volume of the plate was sampled (280 to 310 mL). Subsamples from each fraction were filtered for PON and POC quantification, TEP quantification according to Passow and Alldredge (1995)^61^, microscopic observations, and *nifH* sequencing, *nifH* qPCR as described below. We used the formula described in Riley et al., (2012)^30^ to obtain values associated with fast or slow sinking particles only, quantitatively removing the contribution of other fractions.

### Bottle-net deployment

Vertical 2000-200 m profiles were done using a bottle-net mounted on the CTD rosette frame at stations S03, S04, S05M, S07, S08, S10M, S11 and S12. The bottle-Net consists of a 20-μm conical plankton net housed in a cylindrical PVC pipe^19^. It is lowered with the top cover closed, which is opened at the desired bottom depth (Db, 2000 m) of the tow, remains opened during the ascension of the rosette, and closed again at the upper depth (Du, 200 m) of the water column to be sampled. This results in one integrated sample of 2000 to 200 m per deployment. Once on deck, the bottle-nets were gently rinsed with filtered seawater before retrieving the sample from the collector. At each station, samples were split into aliquots that were processed for were analyzed for *nifH* qPCR. Sampled volume was estimated as the product between the cross-sectional area of the mouth of the bottle-Net (7.5 cm, aspect ratio of 4) and the vertical distance covered by the device from the start of the ascension to the closure of the top cover (1800 m). Blank casts were performed with the bottle-net closed during the entire cast to assess for potential contaminations, and blanks were subtracted.

### *nifH* gene sequencing and bioinformatics

The *nifH* gene was sequenced from a total of 71 samples: 18 samples collected from sediment traps deployed at 170 m, 270 m and 1000 m at S05M and S10M, and 53 samples collected from the suspended, slow sinking and fast sinking fractions of particles collected with the MSC. To that end, DNA was extracted using the DNeasy Plant Mini Kit (Qiagen, Courtaboeuf, France) with additional freeze-thaw bead beating and proteinase K steps before the kit purification^62^. Triplicate nested PCR reactions were conducted using degenerate *nifH* primers^63^. The PCR mix was composed of 5 μL of 5X MyTaq red PCR buffer (Bioline), 1.25 μL of 25 mM MgCl_2_, 0.5 μl of 20 μM forward and reverse primers, 0.25 μL Platinum Taq and 5 μL of DNA extract (1 μL on second round). The reaction volume was adjusted to 25 μL with PCR grade water. Triplicate PCR products were pooled and purified using the Geneclean Turbo kit (MP Biomedicals). Partial adapters were added by ligation at the sequencing facility (Genewiz) and llumina MiSeq 2×300 paired end sequenced. Demultiplexed paired-end sequences were dereplicated, denoised, assembled and chimeras discarded using the DADA2 pipeline^64^. Fifty-thousand to 60,000 reads were obtained per sample. In total, >2.9 millions of high quality *nifH* sequences were obtained resulting in 14925 ASVs (submitted to NCBI with accession numbers SAMN19796776-SAMN19796835). ASVs were annotated down to the genus level using a DADA2 formatted *nifH* gene database (https://github.com/moyn413/nifHdada2). Sequences were grouped into 17 genus according to the database, and sequences not identified to the genus level were grouped as “others”. The *nifH* gene was successfully amplified from all samples.

### Abundance of diazotrophs et contribution to N export fluxes

The abundance of diazotrophs was determined using TaqMan qPCR assays and previously published primer-probe sets for *Trichodesmium*, UCYN-A1, UCYN-B, UCYN-C and γ-24474A11 (the latter hereafter referred to as Gamma-A) targeting the *nifH* gene^65,66,67^. The qPCR was run in 25 µL reactions consisting of 12.25 µL TaqMan PCR Master Mix (Applied Biosystems, Villebon Sur Yvette, France), 1 µL of the forward and reverse primers at 10 µM (HPLC purified, Eurofins, Nantes, France), 0.25 µL probe at 10 µM, 8.25 µL PCR grade water, 0.25 µL bovine serum albumin at 10.08 µg µL^-1^, and 2 µL standard or template sample. The qPCR program was run on a CFX96 Real-Time System thermal cycler (BioRad, Marnes-la-Coquette, France) and consisted of 2 min at 50°C, 10 min at 95°C continued by 45 cycles of 15 s at 95°C and 1 min at 64°C. The annealing temperature was changed to 60°C for UCYN-A1 qPCR runs^67^. Standard dilutions (10^7^-10^1^ gene copies) were run in duplicate, and samples and no-template controls (NTCs) in triplicate. NTCs did not show any amplification. The efficiency was 98-113%. Inhibition tests were carried out on all samples and each primer-probe set by adding the 2 µL of the 10^5^ copies standard to each sample. No inhibition was observed. The limit of detection and detected but not quantifiable limits were 1 and 8 gene copies per reaction, respectively.

The diazotroph turnover rate representing the fraction of surface diazotrophs exported out of the photic layer per day was calculated as follows: Turnover rate (d^-1^) = export flux/abundance in the photic layer. The proportion of PON collected by sediment traps attributed to diazotrophs was calculated based on estimated PON content per cell. For nanoplanktonic UCYN, cell dimensions were directly measured and their biovolume calculated^68^. The C content per cell was estimated from the biovolume according to Verity et al. (1992)^69^ and the N content calculated based on C:N ratios of 5 for UCYN-B ^70,71^ and 8.5 for UCYN-C^50^. For the UCYN-A1-host symbiosis, we considered a size of 1 µm for the UCYN-1 and 2 µm for the host and a C:N ratio of 6.3^72^. For *Trichodesmium*, we considered the average value of 10 ng N per *Trichodesmium* filament from the same region^73^. To circumvent the polyploidy of *Trichodesmium*^74^, we divided the qPCR-based abundances by 12. This ratio was determined for this study by dividing the qPCR-based abundances by the number of *Trichodesmium* filaments counted at S05M and S10M stations at 6 depths in the photic layer (we considered 100 cells per filament of *Trichodesmium*).

### Particle imaging

Seawater samples from traps, MSC and bottle-nets were gently filtered on 0.2 µm (for scanning electron microscopy, SEM) and 2 µm polycarbonate filters (for epifluorescence microscopy) at very low pressure to preserve the particle structure. For epifluorescence microscopy, filters were fixed with paraformaldehyde (2% prepared in filtered seawater) for 10 minutes at ambient temperature and stored at-80°C until visualized using a Zeiss Axioplan (Zeiss, Jena, Germany) microscope fitted with a green (510–560 nm) excitation filter, which targeted the phycoerythrin-rich cells. For SEM, samples were fixed with 2.5% glutaraldehyde and 1.6% PFA for 1 h at room temperature. Subsequently, the filters were rinsed twice in 0.2 µm filtered in seawater during 15 min, rinsed in osmium during 30 min, and rinsed thrice in filtered seawater to eliminate excess osmium. Next, the filters passed through a series of ethanol drying solutions (50, 70, 95 and 100%, 10 min each), and a series of HDMS solutions (30, 50, 80 and 100%, 10 min each). Finally, the filters were air-dried and stored at room temperature until SEM analyses. The filters were visualized onshore using a Phenom-Pro benchtop scanning electron microscope at 10 kV.

### *Tara* Oceans sampling and reads recruitments in metagenomes

23 metagenomes, collected from five stations along the *Tara* Oceans expedition transect and corresponding to a subset of the data presented in (Karlusich JJP, *et al*. 2020), were selected for this study since these stations were the only ones for which : i) diazotrophs were present in surface waters and ii) metagenomes samples from both surface (5m) and mesopelagic waters were collected and sequenced (200-1000 m ; Supplementary Table S2). Briefly, the plankton were separated into discrete organismal size fractions using a serial filtration system^75^ corresponding to picoplankton size (0.2-3 µm), nanoplankton (0.8-3 µm or 0.8-5 µm) and a ‘bulk’ size fraction corresponding to the fraction >0.8 µm or >3 µm. Metagenomes were sequenced as Illumina overlapping paired reads of 100-108 bp, which were merged and trimmed based on quality, resulting in 100-215 bp fragments^76^. Metagenomic reads were recruited against a database of 9 representative genomes of cyanobacterial diazotrophs (Table S1) using BLASTN (v2.9.0; (Altschul *et al*., 1990^77^) with default parameters but limiting the results to one target sequence (--max_target_seqs 1) and keeping only results with an E-value below 1e-30 (-evalue 1e-30) and with a query coverage of, at least, 90% of the read length (-qcov_hsp_perc 90). Following the criteria proposed by Caro-Quintero & Konstantinidis^78^ for metagenome-based prokaryotic genome classification, reads mapping to diazotroph genomes with a percentage of nucleotide identity equal or greater than 95% were taxonomically assigned to each diazotroph genome according to their best hit, except reads that mapped to the ribosomal operon of any of the diazotroph genomes that were filtered out. The number of reads recruited by each diazotroph genome was normalized to the sequencing depth of each sample (Table S2). For Fig. 8, only samples for which genome coverage was higher than 1% were taken into account for each organism.

## Acknowledgements

This research is a contribution of the TONGA project (Shallow hydroThermal sOurces of trace elemeNts: potential impacts on biological productivity and the bioloGicAl carbon pump; TONGA cruise DOI: 10.17600/18000884) funded by the Agence Nationale de la Recherche (grant TONGA ANR-18-CE01-0016 and grant CINNAMON ANR-17-CE2-0014-01), the LEFE-CyBER program (CNRS-INSU), the A-Midex foundation, the Institut de Recherche pour le Développement (IRD). F.M.C-C. and L.G acknowledge funding from the European Union’s Horizon 2020 research programme under the Marie Sklodowska-Curie grant agreement No. 749380 (UCYN2PLAST). The authors warmly thank the crew of the R/V L’Atalante for outstanding shipboard operations. Nagib Bhairy is warmly thanked for his efficient help with MSC deployment and clean CTD rosette management and Vincent Taillandier is tanked for CTD data processing.

## Author contributions

SB designed the experiments and SB, MB and IBF carried them out at sea, with advice from FLM; SB, MB, MC, OG, AT, DS analyzed the samples; FMCC and LG analyzed the *Tara* Oceans metagenomes. SB analyzed the data with the help of MB for the sequencing data. SB prepared the manuscript with contributions of all co-authors.

