## Supplementary material for "Massive export of diazotrophs across the South Pacific tropical Ocean"

**Supplementary information**

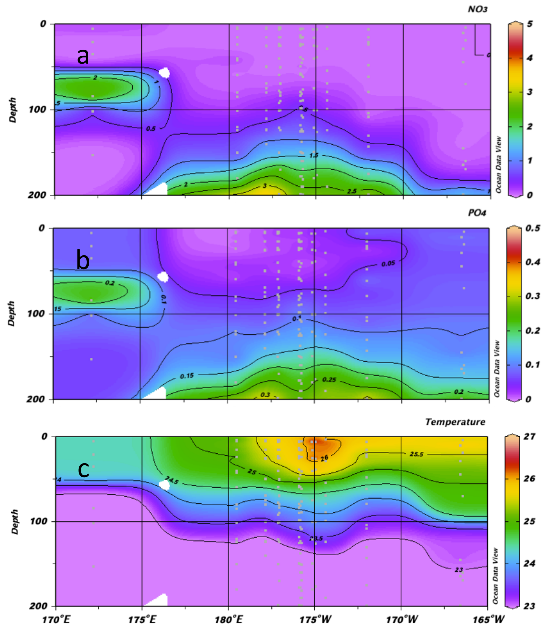


Fig. S1. Horizontal and vertical distributions of a. nitrate concentrations (µmol $L^{-1}$), b. phosphate concentrations (µmol $L^{-1}$) and c. Seawater temperature (°C) across the TONGA transect. Y axis: pressure (dbar), X axis: longitude; grey dots correspond to sampling depths at the various stations.

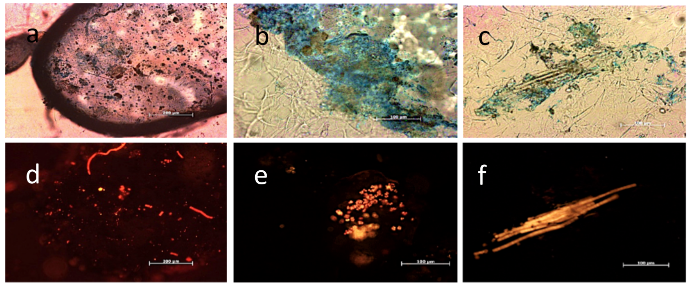


Fig. S2. Microscopy images showing examples of diazotrophs embeded in TEP in sediment trap samples collected at 170 m (a,d), 270 m (b,e), and 1000 m (c-f) at station S10M. Each trio of slides shows an identical image viewed with a-c. white light (upper panels) to highlight staining by Alcian Blue for transparent exopolymeric particles (TEP) seen in blue shades, and d-f. phycoerythrin filter (lower panels). Both panels illustrate the organic matrix surrounding many of the particles in the traps. Visualized on the left panels are phycoerythrin containing cyanobacteria identified as *Trichodesmium* at 1000 m, Crocosphaera-and Synechoccocus like ecotypes at 270 m and a combination of all at 170m. Slides are representative images from these depths. All three traps contained each of the described organisms.

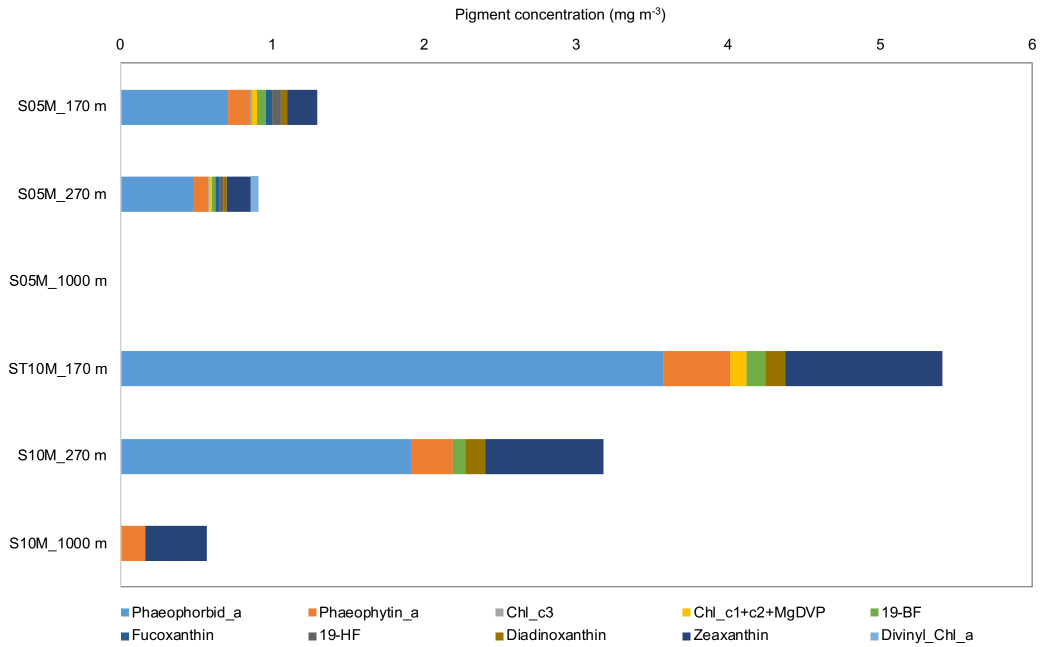


Fig. S3. Pigment concentrations (mg $m^{-3}$) in sediment trap samples at stations S05M and S10M. No data are available at station S05M 1000 m.

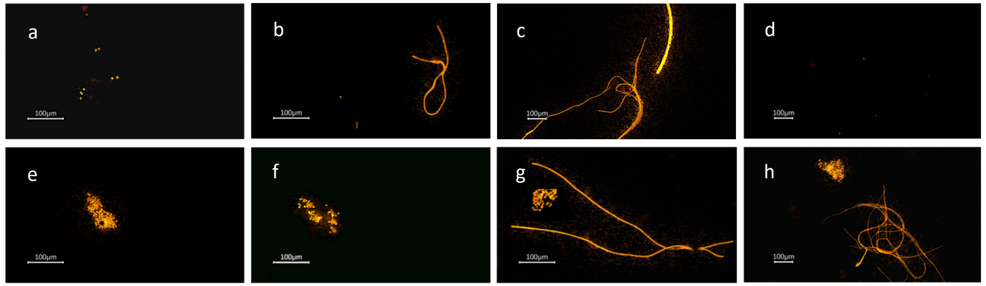


Fig. S4. Epifluorescence microscopy images showing typical examples of phycoerythrin-containing UCYN in the suspended (a-d) and fast sinking (e-h) fractions of the MSC (S05M and S10M, 170 m).

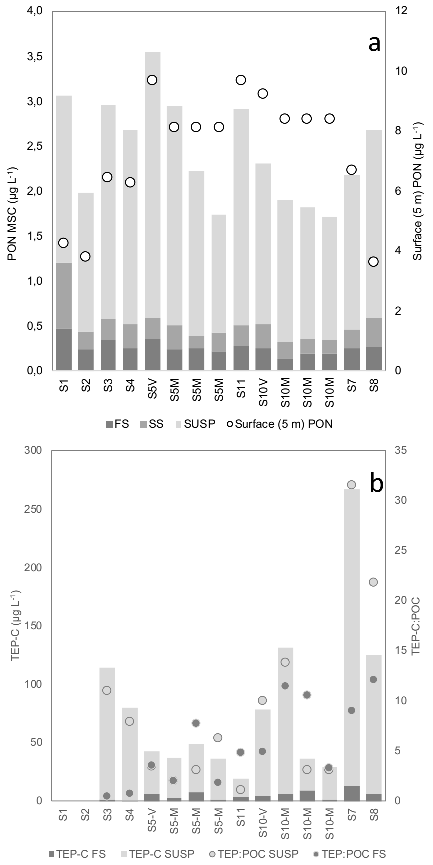


Fig. S5. a. PN concentrations (µg $L^{-1}$) of the suspended, slow and fast sinking pools of the MSC across the transect. Dots represent the surface (5 m) PN pool. b. TEP-C concentrations (µg $L^{-1}$) in the suspended and fast sinking pools. Dots represent the TEP-C:POC ratio for both fractions (suspended: light grey, and fast sinking: dark grey).

Table S1. Contribution of *Trichodesmium* versus UCYN to particulate nitrogen (PN) export fluxes attributted to diazotrophs at stations S05M and S10M at 170 m, 270 m and 1000 m.

| | Station/Depth (m) | 170 m | 270 m | 1000 m |
|---|---|---|---|---|
| Trichodesmium contribution (%) | S05M | 32 | 52 | 80 |
| UCYN contribution (%) | | 68 | 48 | 20 |
| Trichodesmium contribution (%) | S10M | 2 | 8 | 67 |
| UCYN contribution (%) | | 98 | 92 | 33 |

Table S2. Diazotroph genomes used as reference for metagenomic recruitement

| Organism | Genome Accession | References |
|---|---|---|
| Crocosphaera watsonii WH8502 | CAQK01000001 - CAQK01000869 | https://doi.org/10.1111/jpy.12090 |
| Richelia_TARA_PON_109_MAG_00086 | https://figshare.com/articles/dataset/Marine_diazotrophs/14248283 | https://doi.org/10.1101/2021.03.24.436778 |
| Trichodesmium thiebautii H9-4 | SAMN03421272 | https://doi.org/10.1073/pnas.1422332112 |
| Trichodesmium erythraeum IMS101 | SAMN02598485 | https://doi.org/10.1073/pnas.1422332112 |
| Trichodesmium_TARA_AON_82_MAG_00128 | https://figshare.com/articles/dataset/Marine_diazotrophs/14248283 | https://doi.org/10.1101/2021.03.24.436778 |
| Trichodesmium_TARA_IOS_50_MAG_00050 | https://figshare.com/articles/dataset/Marine_diazotrophs/14248283 | https://doi.org/10.1101/2021.03.24.436778 |
| Trichodesmium_TARA_PON_109_MAG_00034 | https://figshare.com/articles/dataset/Marine_diazotrophs/14248283 | https://doi.org/10.1101/2021.03.24.436778 |
| UCYN-A1 | NC_013771.1 | https://doi.org/10.1038/nature08786 |
| UCYN-A2 | JPSP01000000 | https://doi.org/10.1038/ismej.2014.167 |
